# Body mass variations relate to fractionated functional brain hierarchies

**DOI:** 10.1101/2020.08.07.241794

**Authors:** Bo-yong Park, Hyunjin Park, Filip Morys, Mansu Kim, Kyoungseob Byeon, Hyebin Lee, Se-Hong Kim, Sofie Valk, Alain Dagher, Boris C. Bernhardt

**Affiliations:** McConnell Brain Imaging Centre, Montreal Neurological Institute and Hospital, McGill University, Montreal, QC, Canada; School of Electronic and Electrical Engineering, Sungkyunkwan University, Suwon, South Korea; Center for Neuroscience Imaging Research, Institute for Basic Science, Suwon, South Korea; Department of Biostatistics, Epidemiology, and Informatics, University of Pennsylvania, Philadelphia, United States of America; Department of Electrical and Computer Engineering, Sungkyunkwan University, Suwon, South Korea; Department of Family Medicine, St. Vincent’s Hospital, Catholic University College of Medicine, Suwon, South Korea; Otto Hahn Research Group for Cognitive Neurogenetics, Max Planck Institute for Cognitive and Brain Sciences, Leipzig, Germany

**Keywords:** body mass index, functional connectome, manifold, transcriptomic analysis

## Abstract

Variations in body mass index (BMI) have been suggested to relate to atypical brain organization, yet connectome-level substrates of BMI and their neurobiological underpinnings remain unclear. Studying 325 healthy young adults, we examined association between functional connectome organization and BMI variations. We capitalized on connectome manifold learning techniques, which represent macroscale functional connectivity patterns along continuous hierarchical axes that dissociate low level and higher order brain systems. We observed an increased differentiation between unimodal and heteromodal association networks in individuals with higher BMI, indicative of an increasingly segregated modular architecture and a disruption in the hierarchical integration of different brain system. Transcriptomic decoding and subsequent gene enrichment analyses identified genes previously implicated in genome-wide associations to BMI and specific cortical, striatal, and cerebellar cell types. These findings provide novel insights for functional connectome substrates of BMI variations in healthy young adults and point to potential molecular associations.

## Introduction

A high body mass index (BMI) has been recognized as one of the most significant contributors to adverse health and psychological outcomes (Blüher, 2019; James, 2008; World Health Organization, 2020). High BMI is an indicator of obesity, a condition with increasing prevalence worldwide (World Health Organization, 2020) and a critical factor in the development of type 2 diabetes, cardiovascular disease, stroke, cancer, and metabolic syndrome (Jensen et al., 2014; Malik et al., 2013; Raji et al., 2010; Val-Laillet et al., 2015). In addition, multiple neurobiological processes related to obesity have been recognized, including mechanisms regulating eating behaviors, together with genetic and transcriptomic underpinnings (Locke et al., 2015; Martin et al., 2010; Murray et al., 2014; Van Opstal et al., 2018; Steward et al., 2019a; Vainik et al., 2013, 2018; Val-Laillet et al., 2015; Verdejo-Román et al., 2017; Ziauddeen et al., 2015).

Neuroimaging techniques, particularly magnetic resonance imaging (MRI), can identify cerebral substrates associated with BMI by tapping into whole-brain structure, function, and connectivity. Prior structural MRI research has shown that measures of cortical and subcortical morphology robustly correlate with BMI variations in healthy (Herrmann et al., 2019; Marqués-Iturria et al., 2013; Shott et al., 2015; Vainik et al., 2018) and diseased samples (King et al., 2018; Olivo et al., 2017). Multiple task-based functional MRI studies have also shown associations between BMI and brain activations in impulse control and reward processing paradigms (Brooks et al., 2013; Goldstone et al., 2009; Gupta et al., 2018; Van Meer et al., 2019; Opel et al., 2015; Park et al., 2017; Steward et al., 2019b; Stoeckel et al., 2008). On the other hand, functional signatures of BMI at macroscale during resting conditions remain underexplored. Indeed, despite reports exploring associations between BMI and the connectivity of specific regions (García-García et al., 2015; Lips et al., 2014; Park et al., 2015) and larger networks (García-García et al., 2013; Park et al., 2016), whole-brain functional network configurations associated with BMI are less well established. We aim to close this gap in the current work by applying connectome manifold learning techniques to identify functional substrates of BMI in a large population of healthy adults. One appealing feature of these techniques is that they compress high-dimensional connectomes into sets of lower-dimensional eigenvectors that visualize principles of inter-regional connectivity (Margulies et al., 2016). These maps specifically depict the differentiation of brain networks in a continuous manner and index the balance of integration and segregation. These techniques have thus increasingly complemented modular descriptions of brain networks, and offer a data-driven perspective on the gradual hierarchical organization of functional systems (Huntenburg et al., 2018; Margulies et al., 2016). A hierarchical perspective is furthermore supported by work showing a close association between functional gradients and main axes of microstructural differentiation in the cortex, which concomitantly describe a sensory-fugal pattern (Huntenburg et al., 2017; Paquola et al., 2019a, 2020). Connectome manifold learning has furthermore been applied to study healthy aging (Bethlehem et al., 2020; Lowe et al., 2019) and in the hierarchical organization of functional and structural networks in typical and atypical neurodevelopment (Hong et al., 2019; Paquola et al., 2019b; Park et al., 2020a, 2020b). In the context of BMI, these techniques have still not been applied but could promise to identify whether different patterns of functional network integration and segregation underpin inter-individual body mass variations.

As connectome manifold learning can generate cortical maps capturing large-scale principles of brain network organization and hierarchical differentiation, these features can be readily integrated with other aspects of brain organization. As such, spatial associations between connectome gradients and measures of brain morphology and microstructure can be calculated to query shared and unique effects. Furthermore, neural data that is not per se neuroimaging derived is increasingly represented in MRI reference space. One such repository are *post-mortem* gene expression maps disseminated by the Allen Institute for Brain Science (AIBS) (Arnatkeviciute et al., 2019; Chen et al., 2013; Dougherty et al., 2010; Gorgolewski et al., 2014, 2015; Hawrylycz et al., 2012; Kuleshov et al., 2016). This resource can inform spatial association analyses between imaging-derived findings and gene expression patterns. Coupled with gene enrichment analyses (Dougherty et al., 2010; Park et al., 2020a), these approaches can discover molecular, developmental, and disease related processes, and thus provide additional context for MRI-based findings. Recent studies capitalized on transcriptomic decoding to explore underpinnings of brain imaging findings in both healthy and diseased cohorts (Arnatkeviciute et al., 2019; Bertolero et al., 2019; Jahanshad et al., 2013; Paquola et al., 2019b; Park et al., 2020a; Thompson and Fransson, 2016).

Here, we studied associations between macroscale functional connectome organization and variations in BMI. Our functional network analysis was based on the identification of connectome manifolds, which offer a continuous and low dimensional analytical space to interrogate macroscale brain organization and network hierarchy (Burt et al., 2018; Fulcher et al., 2019; Vos de Wael et al., 2020). Capitalizing on the multimodal human connectome project (HCP) dataset (Van Essen et al., 2013; Glasser et al., 2013), we furthermore explored whether associations between functional manifolds and BMI existed above and beyond structural effects as measured by MRI-based measures of cortical thickness, sulco-gyral folding, and intracortical myelin. To explore neurobiological underpinnings of BMI-related whole-brain connectome changes, we performed spatial association analyses to *post-mortem* gene expression data and carried out gene enrichment analyses.

## Results

We studied 325 young and healthy adults (mean ± SD age = 28.56 ± 3.74 years; 55% female; mean ± SD BMI = 26.30 ± 5.16 kg/m^2^, range 16.65 – 47.76 kg/m^2^) from the S900 release of the HCP (Van Essen et al., 2013). See *Methods* for details on participant selection, image processing, and analysis. Reproducibility was studied in an additional 74 unrelated healthy adults from the HCP S1200 release (mean ± SD age = 28.08 ± 3.90 years; 34% female; mean ± SD BMI = 26.17 ± 4.39 kg/m^2^, range 18.89 – 39.47 kg/m^2^), as well as an independent dataset of healthy adults acquired from the St. Vincent’s Hospital (SVH; n = 36; mean ± SD age = 38.78 ± 10.52 years; 47% female; mean ± SD BMI = 29.38 ± 6.29 kg/m^2^, range 23.15 – 57.13 kg/m^2^).

### Macroscale functional manifolds and their association to BMI

We constructed functional connectomes in individual subjects based on correlation analysis of resting-state functional MRI (rs-fMRI) data and estimated functional manifolds (Margulies et al., 2016) using a diffusion map embedding algorithm (Coifman and Lafon, 2006) implemented in the BrainSpace toolbox (https://github.com/MICA-MNI/BrainSpace; see *Methods*) (Vos de Wael et al., 2020). The template manifold was estimated using the group averaged functional connectome, and we aligned individual manifolds to this template using Procrustes rotations (Langs et al., 2015; Vos de Wael et al., 2020). We selected three eigenvectors (M1, M2, M3), explaining approximately 48% of variance in the template affinity matrix (**Fig. 1A–B**). Each eigenvector (also referred to as *gradient*) represents an axis of spatial variation in the functional connectome. In accordance to prior findings in the HCP dataset (Margulies et al., 2016; Vos de Wael et al., 2020), the eigenvectors differentiated primary sensory areas from higher order transmodal areas (M1), visual from somatomotor cortices (M2), and the multiple demand network from task negative systems (M3).

**Fig. 1.**
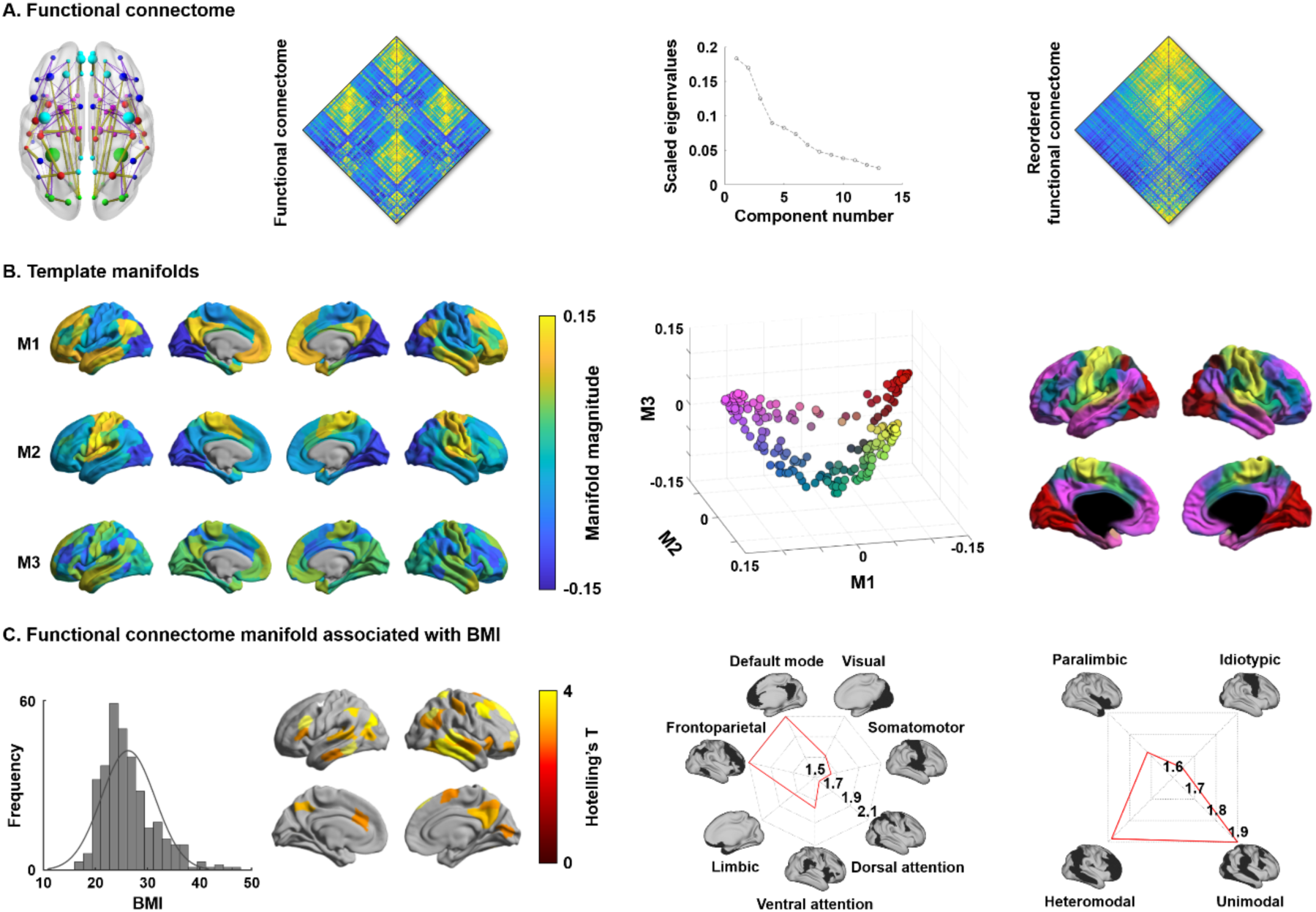
Functional connectome manifolds. **(A)** Functional connectivity schema, a group averaged functional connectome, and a scree plot describing connectome variance across functional components. The reordered functional connectivity matrix according to the first eigenvector (*i.e.*, M1) is shown on the right side. **(B)** Template manifolds built by three dominant eigenvectors (M1, M2, M3) based on the group averaged functional connectome. The scatter plot represents each brain region projected onto the three-dimensional manifold space with different colors, also mapped to the cortical surface for visualization. **(C)** The distribution of BMI is reported on the left. Multivariate association between the three eigenvectors and BMI, highlighting regions showing significant associations to BMI. Findings were corrected for multiple comparisons using a false discovery rate (FDR) < 0.05. Effects were stratified according to intrinsic functional communities (Yeo et al., 2011) and levels of cortical hierarchy (Mesulam, 1998) in the radar plots. *Abbreviation:* BMI, body mass index.

Multivariate analysis associated the three functional gradients with inter-individual differences in BMI, controlling for age and sex. Significant associations were identified in higher order transmodal areas (false discovery rate (FDR) < 0.05; **Fig. 1C**). Stratifying effects according to intrinsic functional communities (Yeo et al., 2011) and the Mesulam model of cortical hierarchical laminar differentiation (Mesulam, 1998), we revealed highest effects in default mode and frontoparietal networks situated in both unimodal and heteromodal association cortices.

### Manifold eccentricity and body mass index

To express the three-dimensional functional manifold structure through a single scalar, we computed the Euclidean distance between the center of template manifold and all data points (*i.e.*, cortical regions) in manifold space (henceforth *manifold eccentricity*) for all participants (**Fig. 2A**). Linear correlations between BMI and functional manifold eccentricity of the regions identified from the multivariate association analysis confirmed significant associations (r = 0.16 and p < 0.001; non-parametric permutation tests; **Fig. 2B**). Correlations were repeated across different levels of cortical hierarchy intersected with significantly associated regions to BMI (see *Fig. 1C*), and strongest effects were identified in unimodal (r = 0.19, p < 0.001) and heteromodal association areas (r = 0.13, p = 0.011; **Fig. 2C**). Paralimbic areas showed marginal effects (r = 0.09, p = 0.06). When analyzing correlations across intrinsic functional communities (Yeo et al., 2011), similar effects were identified with significant associations in default mode and frontoparietal networks as well as somatomotor and dorsal attention networks (p < 0.05; **Fig. S1**).

**Fig. 2.**
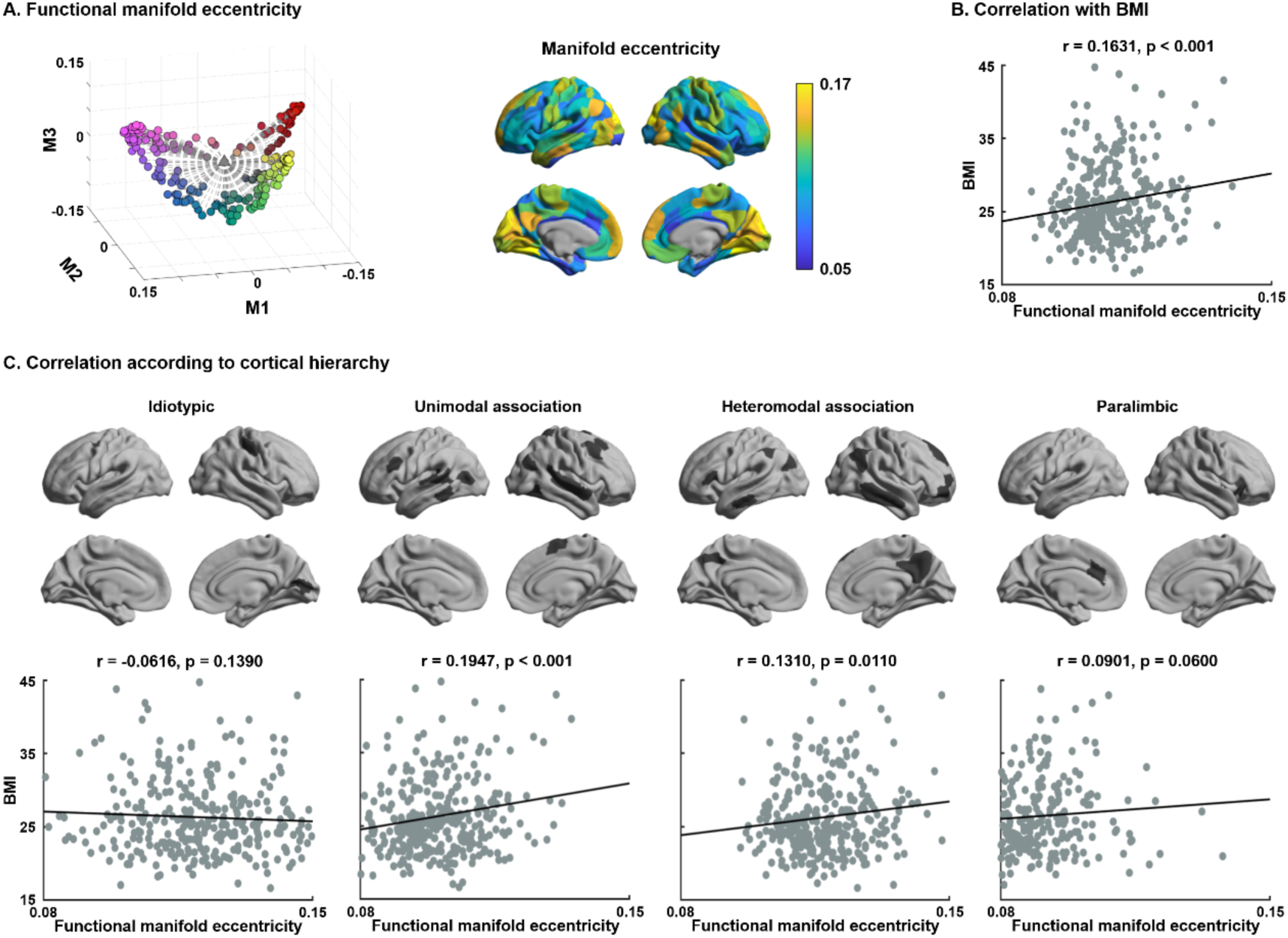
Association between BMI and functional manifold eccentricity. **(A)** Functional manifold eccentricity measured as the Euclidean distance between the center of the template manifold and each data point. **(B)** Linear correlation between BMI and manifold eccentricity. **(C)** Correlations according to a model of cortical hierarchy (Mesulam, 1998). *Abbreviation:* BMI, body mass index.

### Association between manifold eccentricity and modular measures

To assess modular characteristics of BMI-related brain regions in terms of integration and segregation among different functional communities, we calculated within-module degree and participation coefficient (Power et al., 2013; Rubinov and Sporns, 2010) based on modules defined using an established intrinsic functional partitioning (Yeo et al., 2011) (**Fig. 3A–B**). We calculated linear correlation between manifold eccentricity and each modular measure in regions identified in the multivariate analysis (**Fig. 3C**). We found significant positive correlation between manifold eccentricity and within-module degree (r = 0.22, p < 0.001), while participation coefficient showed a negative association (r = −0.16, p = 0.004). These results indicate increased functional segregation of networks in individuals with higher BMI. Similar patterns were observed when defining modules using Louvain community detection algorithm (Blondel et al., 2008) or the Mesulam schema of cortical hierarchy and laminar differentiation (Mesulam, 1998) (**Fig. S2**).

**Fig. 3.**
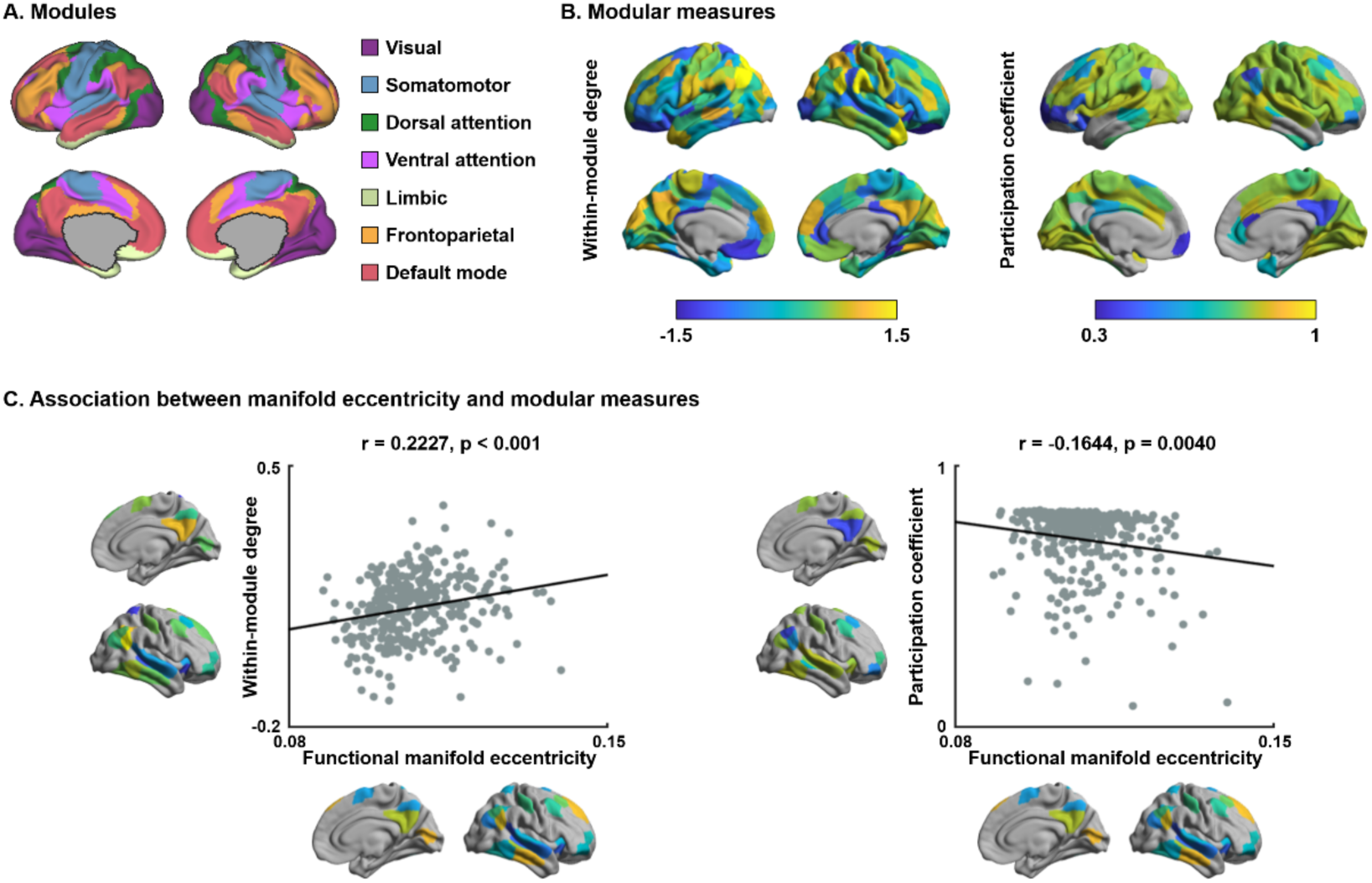
Manifold eccentricity and modular measures. **(A)** Functional communities based on intrinsic functional communities (Yeo et al., 2011). **(B)** Distribution of within-module degree and participation coefficient in the whole-brain. **(C)** Linear correlation between functional manifold eccentricity and modular measures in the identified regions from the multivariate analysis.

### Functional connectome manifold beyond brain structure

Previous studies have reported associations between individual differences in BMI and MRI-based structural indices of cortical thickness, cortical folding, and tissue microstructure (Medic et al., 2016, 2019; Ronan et al., 2019; Vainik et al., 2018; Xu et al., 2013). Here, we explored whether functional connectome manifold findings were, in part, explainable by these underlying structural associations. To this end, we measured cortical morphology (cortical thickness and folding) and intracortical microstructure (the ratio between T1- and T2-weighted imaging contrast, a proxy for intracortical myelin) in the same subjects (**Fig. 4A**) (Glasser and Van Essen, 2011; Glasser et al., 2014; Paquola et al., 2019a). Two analyses were performed. First, we correlated BMI with these indices of brain structure, while controlling for age and sex. While cortical folding was not associated with BMI, a negative effect on cortical thickness was observed in the temporal pole (FDR < 0.05; r = −0.21), and we also found reductions in myelin proxies in occipital, central, and ventrolateral prefrontal regions with increases in BMI (FDR < 0.05; r = −0.35) (**Fig. 4A and S3**). Second, repeating the analysis associating BMI to manifold eccentricity after controlling for the measures of brain structure, findings were consistent in default mode and frontoparietal networks (**Fig. 4B**), while those in limbic networks slightly increased. Collectively, these findings suggest that functional associations were robust above and beyond associations between BMI and cortical (micro)structure.

**Fig. 4.**
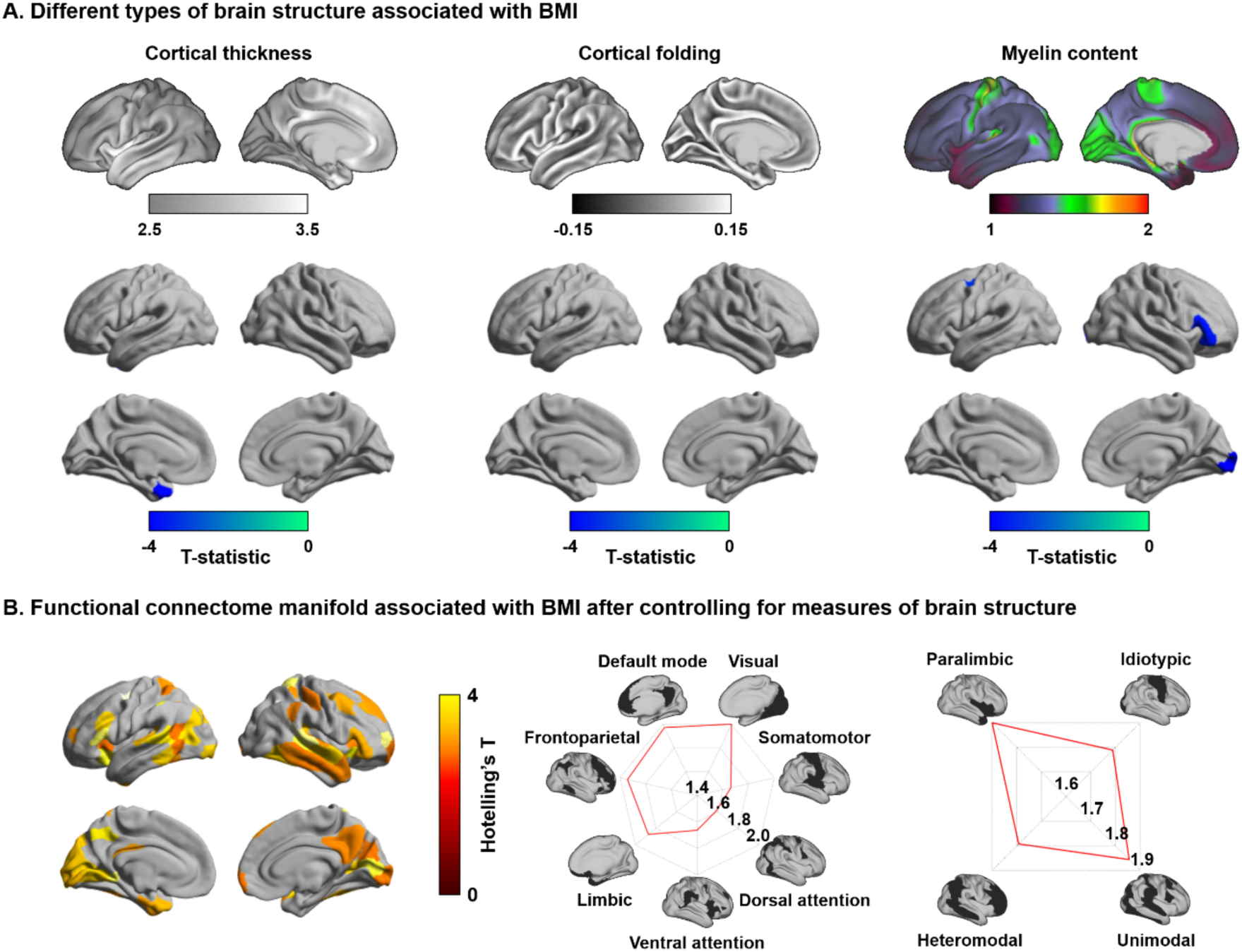
Effects of brain structures. **(A)** Measures of brain structure including cortical morphology and intracortical microstructure. **(B)** Multivariate association of the three manifolds with BMI after controlling for brain structures. *Abbreviation:* BMI, body mass index.

### Transcriptomic association analysis

To provide neurobiological context to our macroscale findings (for details, see *Methods*), we correlated the spatial map of BMI-related functional manifold changes with cortical maps of *post-mortem* gene expression data obtained from the AIBS (Gorgolewski et al., 2014, 2015; Hawrylycz et al., 2012). Among the significantly associated gene lists (FDR < 0.05), only the genes consistently expressed across different donors (FDR < 0.05) (see *Methods*; **Data S1**) (Arnatkeviciute et al., 2019) were fed into the genome-wide association studies using Enrichr (https://amp.pharm.mssm.edu/Enrichr/) (Chen et al., 2013; Kuleshov et al., 2016). These findings pointed to strongest effects for genes previously shown to be associated to BMI (FDR < 0.05; **Fig. 5A**). Further, cell-type specific expression analysis (http://genetics.wustl.edu/jdlab/csea-tool-2/) (Dougherty et al., 2010) suggested that genes associated with BMI-related functional manifold changes were enriched to cortical cells as well as those in striatum and cerebellum (FDR < 0.1; **Fig. 5B**). Specifically, the genes are enriched to the excitatory and inhibitory cells of D1 medium spiny neurons in the striatum and stellate and basket cells in cerebellum, as well as in cortical neurons. These cells are known to indirectly regulate food-related reward processing and appetite (Durst et al., 2019; Matikainen-Ankney and Kravitz, 2018; Timper and Brüning, 2017; Vong et al., 2011).

**Fig. 5.**
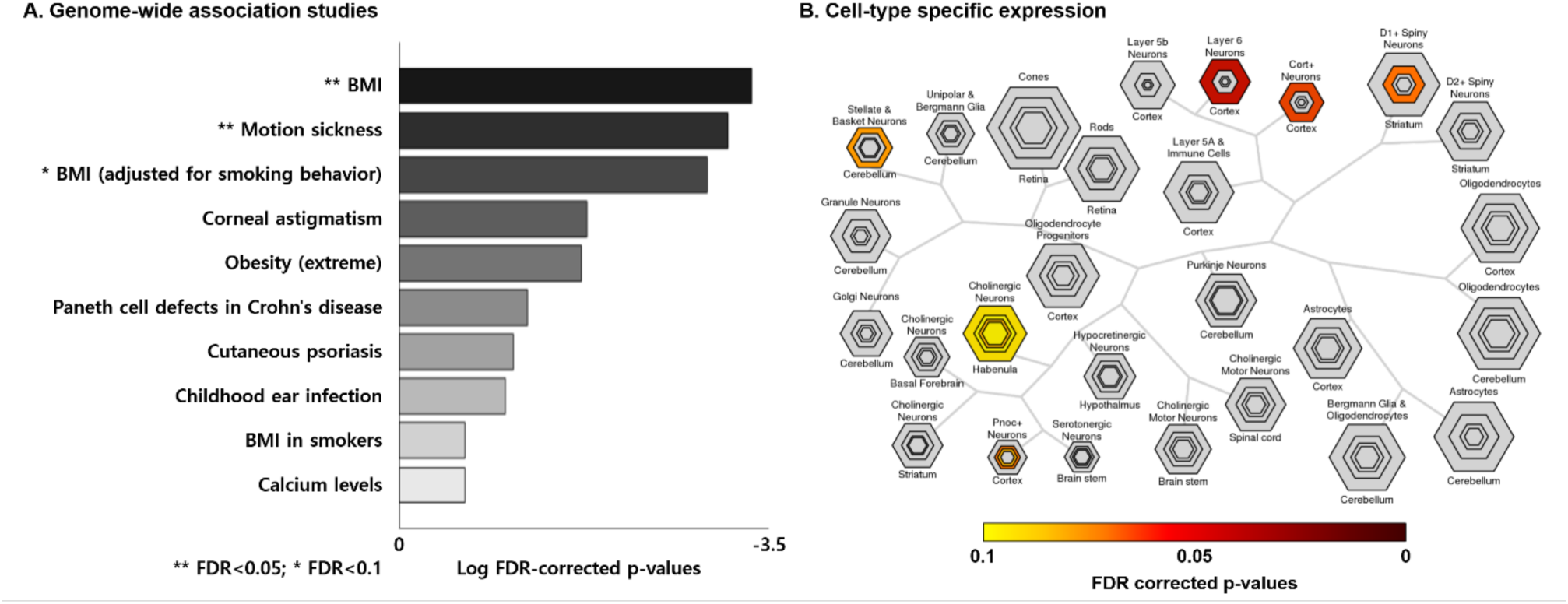
Transcriptomic analysis. **(A)** Top ten categories associated with gene expressions derived from genome-wide association studies. **(B)** Cell-type specific expression analysis identified candidate cell populations associated with genes expressed in the input spatial map (see *Fig. 1C*). *Abbreviations:* BMI, body mass index; FDR, false discovery rate.

### Sensitivity and replication experiments

A series of analyses indicated the robustness of our findings.

#### a) Head motion effect

We repeated the multivariate analyses associating BMI to functional manifold data after controlling for head motion and observed similar spatial patterns (**Fig. S4B**).

#### b) Fluid intelligence and sleep quality effect

It has been shown that BMI also relates to fluid intelligence (Reed et al., 2010; Vainik et al., 2018) and sleep quality (Kohatsu et al., 2006; Vargas et al., 2014). This was also confirmed in this dataset, showing correlations between BMI and fluid intelligence (r = −0.18, p = 0.001) as well as quality of sleep (r = 0.13, p = 0.03). We thus repeated the multivariate association analyses after additionally controlling for these factors. Findings were largely similar to our overall results (**Fig. S4C**).

#### c) Spatial scale

As the main analysis was performed using the Schaefer atlas with 200 parcels (Schaefer et al., 2018), we additionally evaluated the results at both coarser and finer parcellation schemes of 100, 300 and 400 parcels, respectively. Findings were consistent across all parcel resolutions, despite subtle variations in the exact pattern of findings (**Fig. S5, S6**, and **S7**).

#### d) Matrix thresholding

While main findings were based on functional connectomes thresholded at a 10% density as in prior work (Hong et al., 2019; Margulies et al., 2016; Vos de Wael et al., 2020), we also repeated our analysis at 5%, 15%, and 20% densities (**Fig. S8**). We found highly similar patterns at these densities (mean spatial correlation across manifold maps, r = 0.85).

#### e) Group comparison

Instead of carrying out a correlation analysis between functional manifolds and BMI, we also performed a multivariate group comparison analysis to compare cortex-wide manifolds (M1–M3) between individuals with healthy weight (18.5 ≤ BMI < 25) and those with non-healthy weight (BMI ≥ 25). We observed virtually identical results to our main findings (**Fig. S9A**).

#### f) Multivariate association with weight

We additionally performed multivariate association analyses between weight and functional connectome manifolds with controlling for age and sex, as well as height. We found almost unchanged spatial patterns relative to our main findings (**Fig. S9B**).

#### g) Reproducibility in HCP validation dataset

We repeated the main analyses in an independent dataset from the HCP S1200 and found largely consistent results (**Fig. S10**).

#### h) Reproducibility in another dataset

Using an independent dataset with different acquisition parameters (see *Methods*), we replicated our main findings that the functional connectome manifolds in higher order brain networks are associated with BMI (**Fig. S11**).

## Discussion

The human connectome is organized according to multiple processing hierarchies, which allow for integrative and segregated neural functions. Here, we assessed inter-individual differences in this architecture relative to phenotypic variations in body mass index (BMI), an important predictor of health, wellbeing, and life expectancy (Blüher, 2019; James, 2008; World Health Organization, 2020). Our approach leveraged recently established techniques that decompose the whole-brain functional connectome into a set of hierarchical gradients differentiating macroscale systems in a continuous manner along the cortical surface. We observed that unimodal and heteromodal association areas are more differentiated in individuals with higher BMI, suggestive of a potentially disrupted integration between different levels of macroscale hierarchy. Findings remained consistent when additionally controlling for variations in MRI-based measures of cortical morphology and microstructure, suggesting that functional network associations with BMI existed above and beyond regional effects on local brain structure. Functional connectome changes were found in cortical territories known to harbor genes previously implicated in BMI variations, as well as those involved in cortical, striatal, and cerebellar cells. These findings suggest functional network substrates of imbalances in BMI that may ultimately reflect macroscale effects of cellular-genetic associations to BMI.

Manifold learning techniques were utilized to compress and represent high dimensional functional connectomes along a series of spatial gradients. These approaches have recently seen an increasing adoption by the neuroimaging and network neuroscience communities (Burt et al., 2018; Demirtaş et al., 2019; Haak and Beckmann, 2020; Larivière et al., 2019b, 2019a; Müller et al., 2020; Paquola et al., 2019a, 2019b; Park et al., 2020a, 2020b; Vos de Wael et al., 2020; Vos De Wael et al., 2018) to interrogate macroscale neural organization and cortical hierarchy (Hong et al., 2019; Huntenburg et al., 2018; Margulies et al., 2016). Studying the HCP dataset, we identified three functional gradients explaining approximately 50% variance, in agreement with earlier studies in the same dataset (Margulies et al., 2016; Vos de Wael et al., 2020). Notably, associating inter-individual differences in BMI with manifold organization, we observed a marked increase in manifold eccentricity in individuals with higher BMI. These findings were predominantly observed in uni- and heteromodal association cortices that encompass integrative default mode and frontoparietal networks and are reflective of an increased differentiation of these areas to other brain networks. Prior fMRI studies reported atypical intrinsic functional connectivity in individuals with obesity, at both local node and global network levels, relative to individuals with a healthy weight (Chao et al., 2018; Chen et al., 2018; García-García et al., 2013, 2015; Park et al., 2016, 2018). Our findings complement these previous reports focusing on the analysis of connectivity patterns of specific areas (García-García et al., 2013, 2015; Lips et al., 2014; Park et al., 2015) alongside prior graph theoretical analyses (García-García et al., 2015; Park et al., 2016, 2018) in the context of person-to-person variations in BMI. Seed-based and graph theoretical functional connectivity studies found that individuals with obesity showed increased connectivity in nodes belonging to frontoparietal and default mode networks, relative to individuals with healthy weight (Chao et al., 2018; García-García et al., 2013, 2015). These findings are complemented by studies reporting positive associations between overall connectivity degree and broad variability in BMI, again with frequent findings in transmodal areas (Park et al., 2016, 2018). Longitudinal evidence also points to an association between BMI changes and connectivity of reward and frontoparietal networks, both when studying healthy individuals, without any interventions (Park et al., 2019a) and in work that administered repetitive transcranial magnetic stimulation targeting the dorsolateral prefrontal cortex (Kim et al., 2019). Beyond local connectivity, a more recent study reported an increased modular segregation in individuals with obesity compared to individuals with healthy weight, suggesting a shift of brain organization towards more lattice like functional networks as BMI increases (Ottino-González et al., 2020). A more segregated network organization has previously been reported in several psychiatric and neurological diseases, including attention deficit hyperactivity disorder (Cao et al., 2014, 2016, 2013; Liao et al., 2017; Wang et al., 2009), Alzheimer’s disease (Bai et al., 2012; Dai and He, 2014; Liao et al., 2017; Zhao et al., 2012), as well as impulsivity (Davis et al., 2013). These studies noted that increased segregation may reduce global network efficiency and delay information transfer between nodes (Avena-Koenigsberger et al., 2014, 2018, 2019), potentially contributing to cognitive decline (Liao et al., 2017; Sporns, 2013). Based on these studies and our findings, the observed increased segregation of unimodal and heteromodal association cortices in individuals with high BMI could potentially reflect disruptions in feedforward and feedback processing, and potentially indicate atypical cognitive flexibility in individuals with high BMI (Martin et al., 2010; Moore et al., 2017; Moreno-Lopez et al., 2016; Morys et al., 2020; Park et al., 2016; Vainik et al., 2018; Val-Laillet et al., 2015; Whitmer et al., 2007; Ziauddeen et al., 2015). Of note, these findings were largely consistent when incorporating a range of potential confounds, including sleep quality as well as fluid intelligence. Moreover, we could observe similar patterns in an initially held out HCP subsample, as well as in a completely different dataset acquired in South Korea, supporting that our findings are overall robust.

In addition to conforming with established models of hierarchical network organization (Margulies et al., 2016; Mesulam, 1998), another appealing feature of the manifold framework is the ability to project connectome-derived findings back to cortical surfaces. In our analyses, this allowed for an integration of functional findings with morphological and microstructural measures in the same participants. Previous studies have explored morphological substrates of BMI variations, reporting cortical thinning in lateral prefrontal, entorhinal, and parahippocampal regions as BMI increases, indicating that overweight and obese people have reduced cortical thickness compared to people with a normal body weight (Medic et al., 2016; Ronan et al., 2019; Shaw et al., 2018; Vainik et al., 2018; Veit et al., 2014; Westwater et al., 2019). A recent multi-site study confirmed that high BMI (≥ 30) relates to reduced cortical thickness in temporal and frontal regions (Opel et al., 2020). In our study, we observed diffuse tendencies for decreased cortical thickness in individuals with higher BMI, with significant peak effects in temporopolar cortices. Findings were complemented by microstructural associations in primary sensory and ventrolateral prefrontal cortices, potentially indicative of myelin anomalies in individuals with high BMI that have already been suggested based on different methodologies (Metzler-Baddeley et al., 2018; Sena et al., 1985; Xiao et al., 2018). Notably, however, we observed virtually unchanged associations between BMI and functional connectome manifold changes when controlling for MRI-derived indices of morphology and microstructure, indicating that the functional connectome reorganization situated in higher order brain regions occurred above and beyond these underlying structural variations.

In addition to analyses of regional morphology and microstructure, we performed a transcriptomic association analysis based on *post-mortem* gene expression maps provided by the Allen Institute for Brain Sciences (AIBS). Although such transcriptomic associations were thus based on a different dataset, equivalent approaches have been increasingly adopted in neuroimaging research to identify potential molecular patterns that covary with macroscopic findings (Arnatkeviciute et al., 2019; Chen et al., 2013; Dougherty et al., 2010; Gorgolewski et al., 2015; Hawrylycz et al., 2012; Kuleshov et al., 2016). In our work, spatial association analyses pointed to specific gene sets, which we then decoded against findings from previously reported genome-wide association studies. This analysis demonstrated that the spatial pattern of functional connectome manifold changes co-localizes with genes previously implicated in BMI variations; as such, our enrichment analyses highlights that the macroscale functional connectome associations reported here likely reflect genetically mediated processes. Additional gene enrichment analyses furthermore suggested that the identified genes are mainly expressed by cortical neurons, together with cells in the cerebellum as well as D1 medium spiny neurons in the striatum. Although these associations are indirect and based on different samples, they may extend and recapitulate computational theories on circuit mechanisms contributing to BMI, and notably point to an atypical organization of dopaminergic circuits involving mesolimbic as well as cortical control systems.

In sum, the current study identified functional connectome substrates of BMI variations in healthy young adults based on advanced connectome manifold learning. Our findings point to a fractionation in the modular and hierarchical organization of the brain, specifically between unimodal and heteromodal association cortices. These findings were found to be robust across a range of confounds and baseline variations in cortical morphology, and could be replicated in two additional datasets. Further transcriptomic decoding showed that these patterns were associated with genetic factors contributing to BMI variations as well, and pointed to the expressions of cortical as well as subcortical neurons implicated in dopamine signaling. Our findings, thus, provide new insights into coupled macroscale and molecular underpinnings of BMI variations in the adult human brain.

## Methods

### Participants

We obtained the minimally processed imaging and phenotypic data from the S900 release of HCP (Van Essen et al., 2013). We excluded participants who did not complete full imaging data (*i.e.*, T1-weighted, T2-weighted, and rs-fMRI) and who were genetically related (*i.e.*, twin pairs), resulting in a total of 325 participants (mean ± SD age = 28.56 ± 3.74 years; 55% female). The mean BMI of the participants was 26.30 kg/m^2^ with SD of 5.16, range = 16.65 – 47.76 kg/m^2^), and the proportion of underweight (BMI < 18.5 kg/m^2^), healthy weight (18.5 ≤ BMI < 25 kg/m^2^), overweight (25 ≤ BMI < 30), and obesity (BMI ≥ 30) was 6:143:113:63. We additionally obtained data from the S1200 release of HCP to replicate our findings. The same exclusion criteria were applied. A total of 74 participants (mean ± SD age = 28.08 ± 3.90 years; 34% female; mean ± SD BMI = 26.17 ± 4.39 kg/m^2^, range 18.89 – 39.47 kg/m^2^) were enrolled and the ratio of healthy weight, overweight, and obesity was 30:29:15. All MRI data used in this study were publicly available and anonymized. Participant recruitment procedures and informed consent forms, including consent to share de-identified data, were previously approved by the Washington University Institutional Review Board as part of the HCP.

To replicate findings, we collected an independent dataset from an independent site (St. Vincent’s Hospital (SVH): n = 36; mean ± SD age = 38.78 ± 10.52 years; 47% female; mean ± SD BMI = 29.38 ± 6.29 kg/m^2^, range 23.15 – 57.13 kg/m^2^). Data collection and usage were approved from the Institutional Review Boards of the Catholic University of Korea (no. XC15DIMI0012, approved March 2015) and written and informed consent was obtained from all participants.

### MRI acquisition

#### a) HCP

The HCP imaging data were scanned using a Siemens Skyra 3T at Washington University. The T1-weighted images were acquired using a magnetization-prepared rapid gradient echo (MPRAGE) sequence (repetition time (TR) = 2,400 ms; echo time (TE) = 2.14 ms; field of view (FOV) = 224 × 224 mm^2^; voxel size = 0.7 mm^3^; and number of slices = 256). The T2-SPACE sequence was used for scanning T2-weighted structural data, and the acquisition parameters were the same as the T1-weighted data except for the TR (3,200 ms) and TE (565 ms). The rs-fMRI data were collected using a gradient-echo EPI sequence (TR = 720 ms; TE = 33.1 ms; FOV = 208 × 180 mm^2^; voxel size = 2 mm^3^; number of slices = 72; and number of volumes = 1,200). During the rs-fMRI scan, participants were instructed to keep their eyes open looking at a fixation cross. Two sessions of rs-fMRI data were acquired; each of them contained data of left-to-right and right-to-left phase-encoded directions, providing up to four time series per participant.

#### b) SVH

The SVH imaging data were scanned using a Siemens Magnetom 3T scanner equipped with a 32-channel head coil. The T1-weighted images were acquired using a MPRAGE sequence (TR = 1,900 ms; TE = 2.49 ms; FOV = 250 × 250 mm^2^; voxel size = 1 mm^3^; and number of slices = 160). The rs-fMRI data were collected using a gradient-echo EPI sequence (TR = 2,490 ms; TE = 30 ms; FOV = 220 × 220 mm^2^; voxel size = 3.4 × 3.4 × 3 mm^3^; number of slices = 36; and number of volumes = 150).

### Data preprocessing

#### a) HCP

HCP data were minimally preprocessed using FSL, FreeSurfer, and Workbench (Fischl, 2012; Glasser et al., 2013; Jenkinson et al., 2012). Structural MRI data were corrected for gradient nonlinearity and b0 distortions, and co-registration was performed between the T1- and T2-weighted data using a rigid-body transformation. Bias field was adjusted using the inverse intensities from the T1- and T2-weighting. Processed data were nonlinearly registered to MNI152 space, and white and pial surfaces were generated by following the boundaries between different tissues (Dale et al., 1999; Fischl et al., 1999a, 1999b). The midthickness surface was generated by averaging white and pial surfaces, and it was used to generate the inflated surface. The spherical surface was registered to the Conte69 template with 164k vertices (Van Essen et al., 2012) using MSMAll (Glasser et al., 2016) and downsampled to a 32k vertex mesh. The rs-fMRI data were preprocessed as follows: First, EPI distortions and head motion were corrected, and data were registered to the T1-weighted data and subsequently to MNI152 space. Magnetic field bias correction, skull removal, and intensity normalization were performed. Noise components attributed to head movement, white matter, cardiac pulsation, arterial, and large vein related contributions were removed using FMRIB’s ICA-based X-noiseifier (ICA-FIX) (Salimi-Khorshidi et al., 2014). Preprocessed time series were mapped to the standard grayordinate space, with a cortical ribbon-constrained volume-to-surface mapping algorithm. The total mean of the time series of each left-to-right/right-to-left phase-encoded data was subtracted to adjust the discontinuity between the two datasets and they were concatenated to form a single time series data.

#### b) SVH

SVH data were processed using the fusion of the neuroimaging preprocessing (FuNP) pipeline integrating AFNI, FreeSurfer, FSL, and ANTs (Avants et al., 2011; Cox, 1996; Fischl, 2012; Glasser et al., 2013; Jenkinson et al., 2012; Park et al., 2019b). T1-weighted data were processed using the equivalent procedures as in HCP data. The rs-fMRI preprocessing involved removal of the first 10 s (5 volumes) to allow for magnetic field saturation. Head motion and slice timing were corrected, and volumes with frame-wise displacement > 0.5 mm were removed. After removing non-brain tissues, intensity was normalized. Nuisance variables of head motion, white matter, cerebrospinal fluid, cardiac pulsation, and arterial and large vein related contributions were removed using ICA-FIX (Salimi-Khorshidi et al., 2014). Registration from fMRI onto the T1-weighted data and subsequently onto the MNI 3 mm^3^ standard space was performed. Data underwent a band-pass filter with a pass band from 0.009 and 0.08 Hz and spatial smoothing with a full width at half maximum of 5 mm. The processed fMRI data were mapped to the cortical surface with a cortical ribbon-constrained volume-to-surface mapping algorithm.

### Low dimensional functional manifold identification

We generated functional connectomes by computing linear correlations of the time series between two different brain regions, using the Schaefer 7-network based atlas with 200 parcels (Schaefer et al., 2018). Correlation coefficients underwent Fisher’s r-to-z transformations to render data more normally distributed (Thompson and Fransson, 2016). Cortex-wide functional manifolds (*i.e.*, the principal eigenvectors explaining spatial shifts in the functional connectome) were estimated using BrainSpace (https://github.com/MICA-MNI/BrainSpace) (Vos de Wael et al., 2020). First, a template manifold was estimated from a group average functional connectome (**Fig. 1A**). A similarity matrix, capturing similarity of connections among different brain regions, was constructed using a normalized angle kernel with a connection density of 10%. We generated the connectome manifolds (**Fig. 1B**) using diffusion map embedding (Coifman and Lafon, 2006), which is robust to noise and computationally efficient compared to other non-linear manifold learning techniques (Von Luxburg, 2007; Tenenbaum et al., 2000). It is controlled by two parameters α and t, where α controls the influence of the density of sampling points on the manifold (α = 0, maximal influence; α = 1, no influence) and t scales eigenvalues of the diffusion operator. As in prior applications (Hong et al., 2019; Margulies et al., 2016; Paquola et al., 2019a; Vos de Wael et al., 2020), we set α = 0.5 and t = 0 to retain the global relations between data points in the embedded space. In this new manifold, interconnected brain regions are closely located, and the regions with weak inter-connectivity located farther apart. The individual-level manifolds were estimated and aligned to the template manifold via Procrustes alignment (Langs et al., 2015; Vos de Wael et al., 2020).

### Macroscale connectome associated with body mass index

We performed multivariate association analysis between BMI and the first three functional manifolds, which explained approximately 50% in connectome variance, with the model controlling for age and sex. We corrected for multiple comparisons using the FDR procedure (Benjamini and Hochberg, 1995). We summarized multivariate association statistics within well-established resting-state functional communities (Yeo et al., 2011) and with respect to proposed levels of cortical hierarchy (Mesulam, 1998) (**Fig. 1C**). We simplified the multivariate manifolds into a single scalar value by calculating the Euclidean distance between the center of template manifold and all data points (*i.e.*, brain regions) in the manifold space for each individual, which was referred to as manifold eccentricity (**Fig. 2A**) (Bethlehem et al., 2020; Park et al., 2020b). We calculated linear correlation between BMI and the manifold eccentricity of the identified regions derived from the multivariate analysis (**Fig. 2B**). We also stratified associations according to four cortical hierarchy levels (**Fig. 2C**) (Mesulam, 1998) and seven functional communities (**Fig. S1**) (Yeo et al., 2011). The significance of the correlation was assessed using 1,000 permutation tests by randomly shuffling participants. A null distribution was constructed and the real correlation strength was deemed significant if it did not belong to the 95% of the distribution (two-tailed p < 0.05).

### Modular characteristic according to the functional manifold eccentricity

To assess how the modular architecture changes according to the functional connectome manifolds, we calculated Pearson’s correlation between the manifold eccentricity and within-module degree and participation coefficient (Power et al., 2013; Rubinov and Sporns, 2010) in regions identified by the multivariate analysis (**Fig. 3** and **S2**). Modules were defined using established intrinsic functional communities (Yeo et al., 2011), a Louvain community detection algorithm (Blondel et al., 2008), and a schema of cortical hierarchy (Mesulam, 1998). Within-module degree is the degree centrality within a module, indicating the intra-modular connections, while participation coefficient represents inter-modular connections (Power et al., 2013; Rubinov and Sporns, 2010). In other words, high within-module degree represents that a given node has the property of being a hub node within a given module. In contrast, high participation coefficient indicates the node has edges distributed equally to other modules. The significance of the associations between manifold eccentricity and modular measures were assessed using 1,000 permutation tests by randomly shuffling subjects.

### Associations to brain structure

To assess morphological and microstructural underpinnings (**Fig. 4A**), we first correlated BMI with MRI-based measures of cortical morphology, *i.e.*, cortical thickness and cortical folding, and *in vivo* proxies of intracortical microstructure, *i.e.*, the ratio of the T1-weighted and T2-weighted imaging contrast in voxels between the white and pial surfaces (Glasser and Van Essen, 2011; Glasser et al., 2014; Paquola et al., 2019a). We also repeated the association analysis between BMI and manifold changes, after controlling for regional structural indices (**Fig. 4B**).

### Transcriptomic association analysis

Transcriptomic association analysis explored neurobiological underpinnings of functional manifold eccentricity (**Fig. 5**) (Arnatkeviciute et al., 2019; Chen et al., 2013; Dougherty et al., 2010; Gorgolewski et al., 2015; Hawrylycz et al., 2012; Kuleshov et al., 2016). Specifically, we correlated the t-statistical map of manifold changes associated with BMI with *post-mortem* gene expression map from the AIBS using the Neurovault gene decoding tool (Gorgolewski et al., 2015; Hawrylycz et al., 2012). Neurovault implements mixed-effect analysis to estimate associations between the input t-statistic map and the genes of AIBS donor brains yielding the gene symbols associated with the input t-statistic map. Gene symbols that passed a significance level of FDR-corrected p < 0.05 were further tested whether they are consistently expressed across donors using abagen (https://github.com/rmarkello/abagen) (Arnatkeviciute et al., 2019). For each gene, we estimated the whole-brain expression map for each donor, and correlated the maps between all pairs of donors. Genes showing consistent a whole-brain expression pattern across donors (FDR < 0.05) were compared with genes extracted from genome-wide association studies using Enrichr (https://amp.pharm.mssm.edu/Enrichr/) (Chen et al., 2013; Kuleshov et al., 2016). Then we fed the consistent genes into the cell-type specific expression analysis (http://genetics.wustl.edu/jdlab/csea-tool-2/) to identify candidate cell populations likely to be associated with input gene lists (Dougherty et al., 2010). Significances were assessed using a z-score modification of Fisher’s exact test and FDR correction.

### Sensitivity and reproducibility analyses

#### a) Head motion effect

To assess the effects of head motion on functional connectome manifolds, we repeated multivariate association analysis with controlling for age and sex, as well as head motion that calculated based on the frame-wise displacement during fMRI scan (**Fig. S4B**).

#### b) Fluid intelligence and sleep quality effect

It is known that BMI is related to fluid intelligence (Reed et al., 2010; Vainik et al., 2018) and sleep quality (Kohatsu et al., 2006; Vargas et al., 2014). To assess the relationship between BMI and these factors, we obtained fluid intelligence score measured using Penn Progressive Matrices (Bilker et al., 2012) and quality of sleep measured using Pittsburgh Sleep Quality Index (Backhaus et al., 2002; Buysse et al., 1989; Carpenter and Andrykowski, 1998). We repeated multivariate analyses to associate functional connectome manifolds and BMI with controlling for age and sex, as well as fluid intelligence and sleep quality (**Fig. S4C**).

#### c) Spatial scale

To evaluate the impact of spatial scale, we repeated our analyses across different scales of the Schaefer atlas (*i.e.*, 100, 300, or 400 regions) (**Fig. S5, S6, and S7**) (Schaefer et al., 2018).

#### d) Matrix thresholding

We repeated functional manifold estimation using functional connectomes with different levels of density from 5% to 20% with an interval of 5% (**Fig. S8**).

#### e) Group comparison

We compared functional connectome manifolds spanned by M1–M3 between individuals with healthy weight (18.5 ≤ BMI < 25) and non-healthy weight (BMI ≥ 25), controlling for age and sex, to assess whether the findings from multivariate association to BMI are similar to those from multivariate group comparison (**Fig. S9A**). The six underweight (BMI < 18.5) individuals were excluded.

#### f) Multivariate association with weight

We performed multivariate analyses to associate functional connectome manifolds with weight after controlling for age, sex, and height to additionally validate our main findings (**Fig. S9B**).

#### g) Reproducibility in HCP validation dataset

We performed the same analyses using the validation dataset obtained from the S1200 release of the HCP to replicate our findings (n = 74) (**Fig. S10**).

#### h) Reproducibility in another dataset

We replicated our findings using the independent SVH dataset (n = 36) (**Fig. S11**).

## Acknowledgments

Data were provided, in part, by the Human Connectome Project, WU-Minn Consortium (Principal Investigators: David Van Essen and Kamil Ugurbil; 1U54MH091657) funded by the 16 NIH Institutes and Centers that support the NIH Blueprint for Neuroscience Research, and by the McDonnell Center for Systems Neuroscience at Washington University. Dr Bo-yong Park was funded by Molson Neuro-Engineering fellowship by Montreal Neurological Institute and Hospital (MNI) and the Fonds de la Recherche due Québec – Santé (FRQ-S). Drs Bo-yong Park and Boris C. Bernhardt are jointly funded through an MNI-Cambridge collaborative award. Dr Boris Bernhardt further acknowledges research support from the National Science and Engineering Research Council of Canada (NSERC Discovery-1304413), the Canadian Institutes of Health Research (CIHR FDN-154298), SickKids Foundation (NI17-039), Azrieli Center for Autism Research (ACAR-TACC), and the Tier-2 Canada Research Chairs program.

## Author contributions

B.P. and B.C.B. designed the experiments, analyzed the data, and wrote the manuscript. H.P., K.B., H.L., and S.H.K. aided the experiments. H.P., F.M., M.K., S.V., and A.D. reviewed the manuscript. B.P. and B.C.B. are the corresponding authors of this work and have responsibility for the integrity of the data analysis.

---

**Fig. S1.**
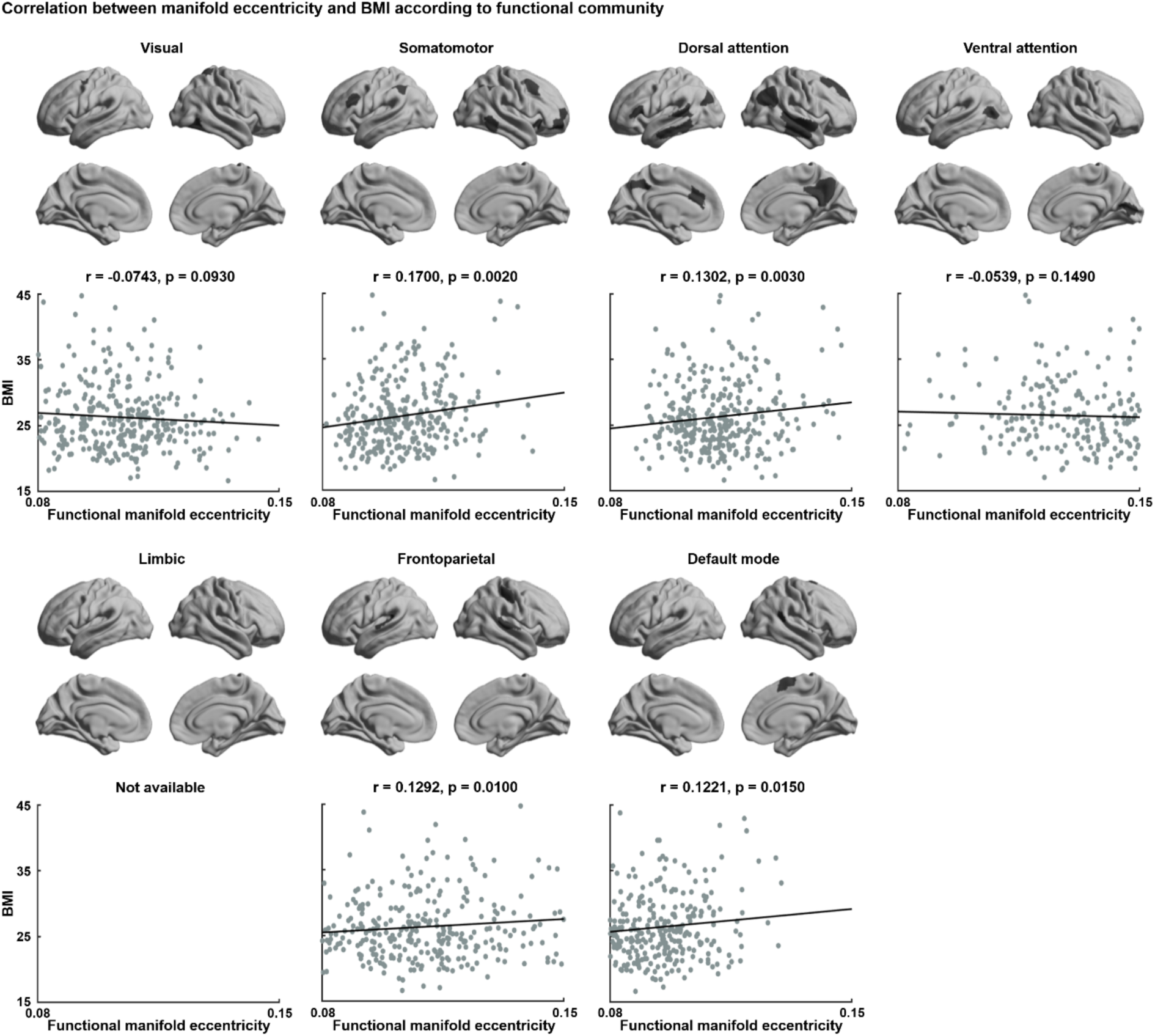
Association between BMI and functional manifold eccentricity. Linear correlations between BMI and functional manifold eccentricity according to intrinsic functional communities (Yeo et al., 2011).

**Fig. S2.**
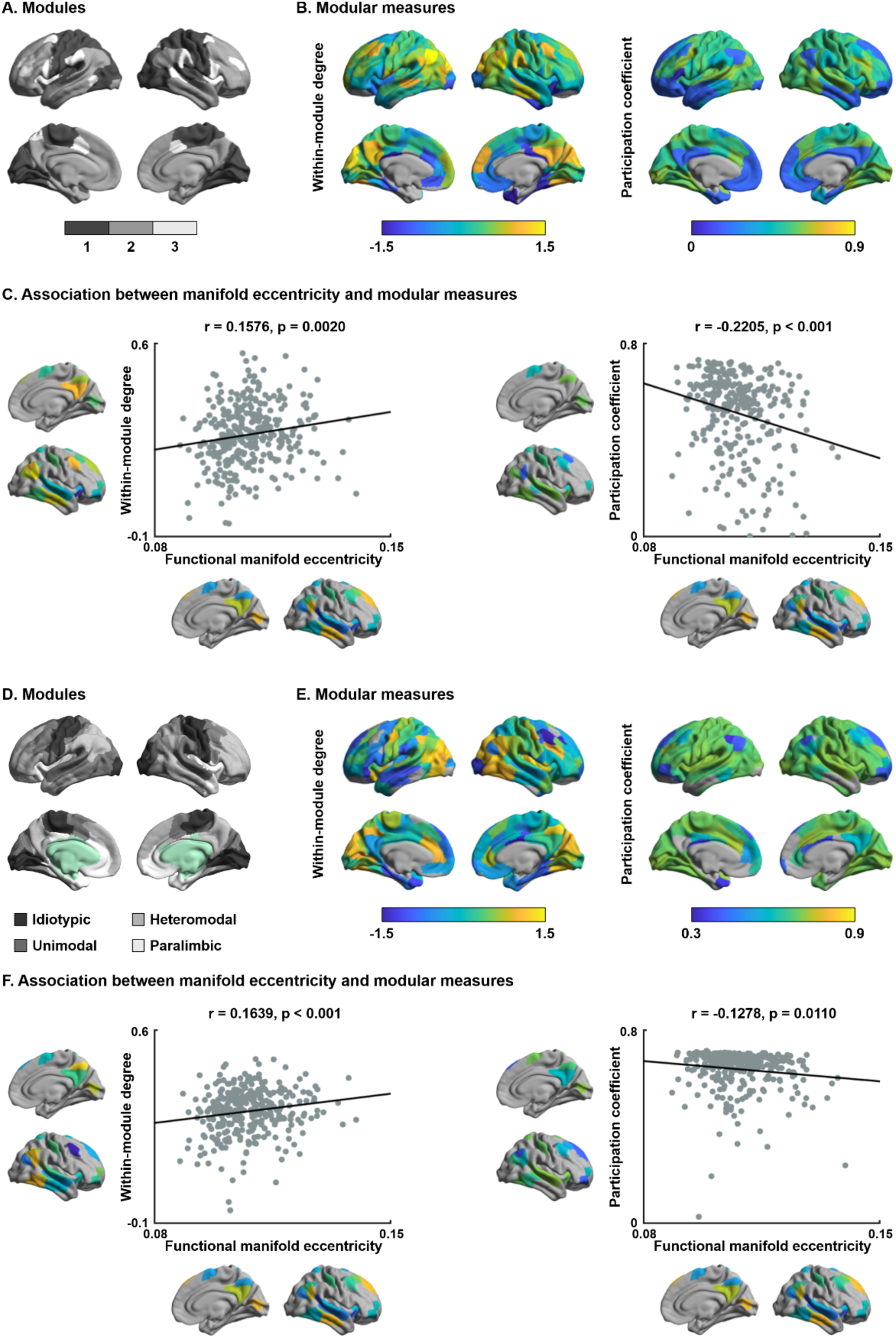
Manifold eccentricity and modular measures. (**A–C**) Modules defined using Louvain community detection (Blondel et al., 2008) and (**D–F**) cortical hierarchy (Mesulam, 1998). For details, see *Fig. 3*.

**Fig. S3.**
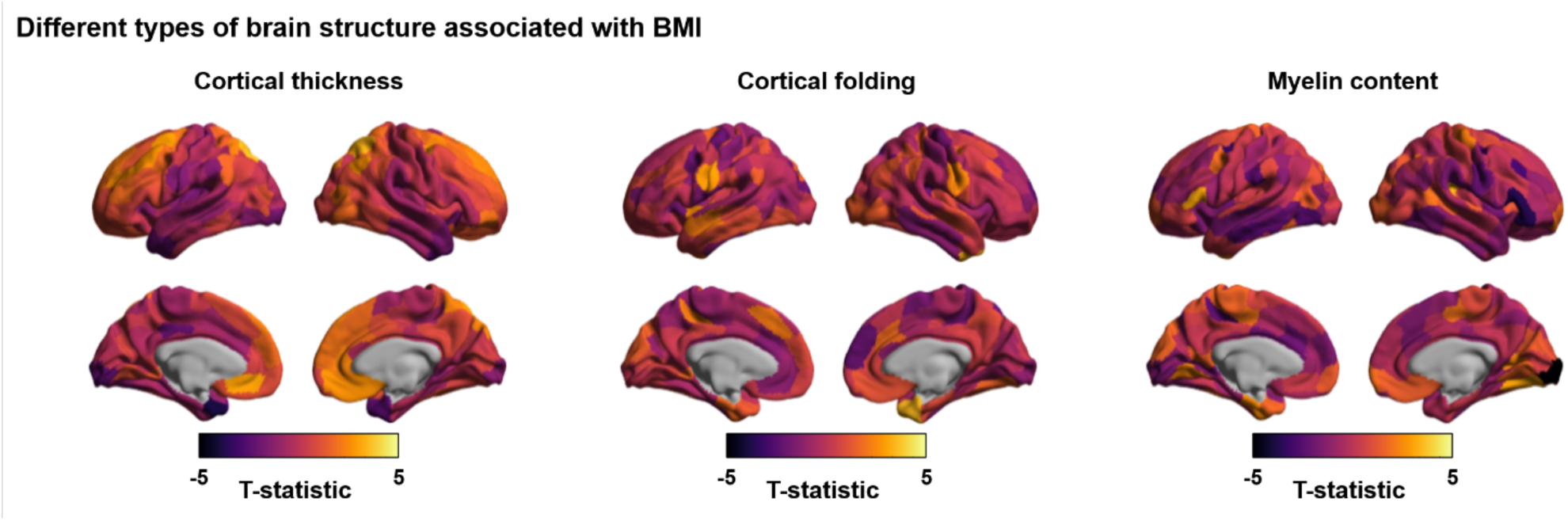
Effects of brain structures. The t-statistics of the association between brain structures (*i.e.*, cortical thickness, cortical folding, and myelin content) and BMI.

**Fig. S4.**
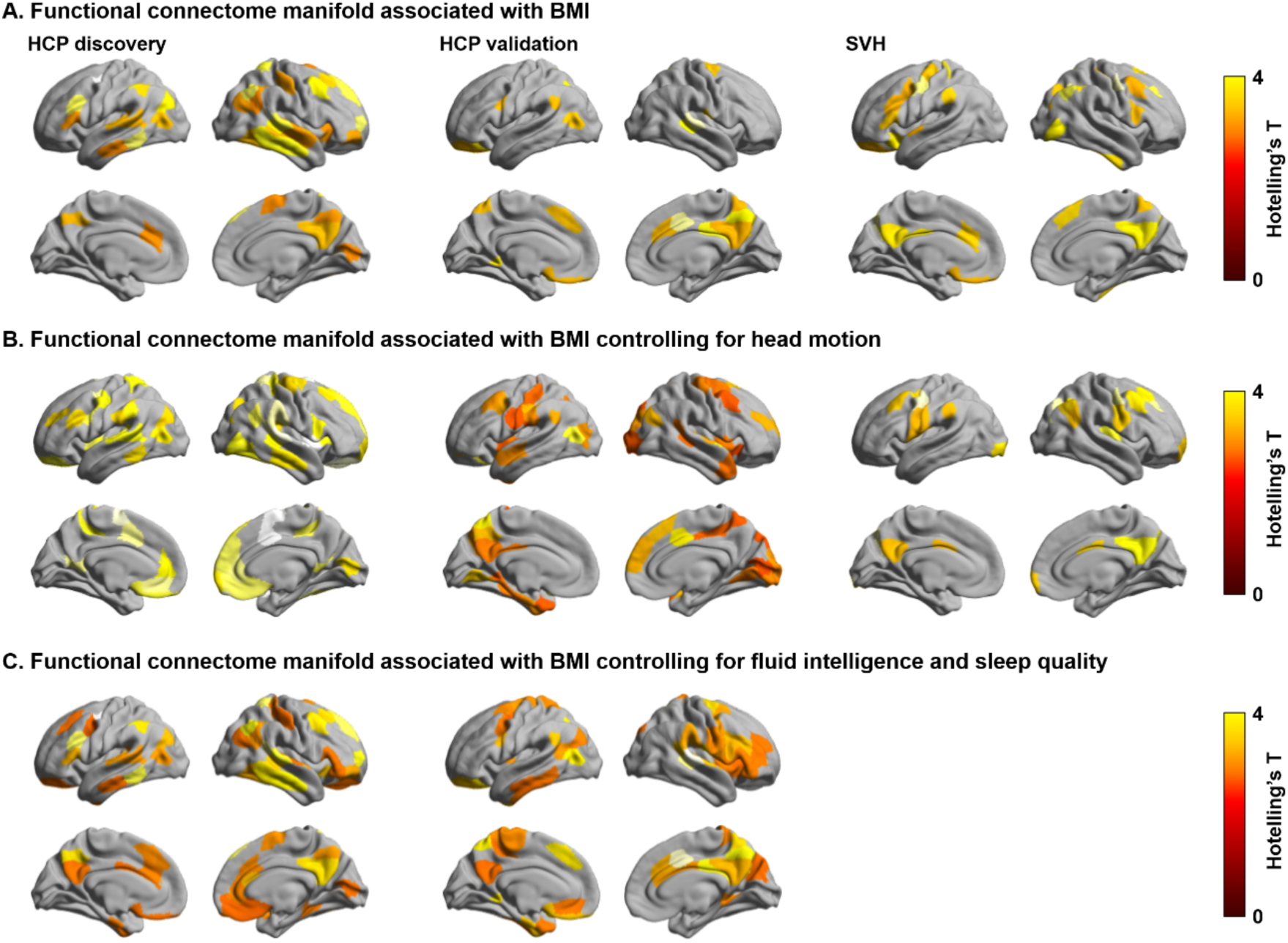
Head motion effect. **(A)** Multivariate association between the three manifolds and BMI without and **(B)** with controlling for head motion. **(C)** Multivariate association results after controlling for fluid intelligence and sleep quality. *Abbreviations:* BMI, body mass index; HCP, human connectome project; SVH, St. Vincent’s Hospital.

**Fig. S5.**
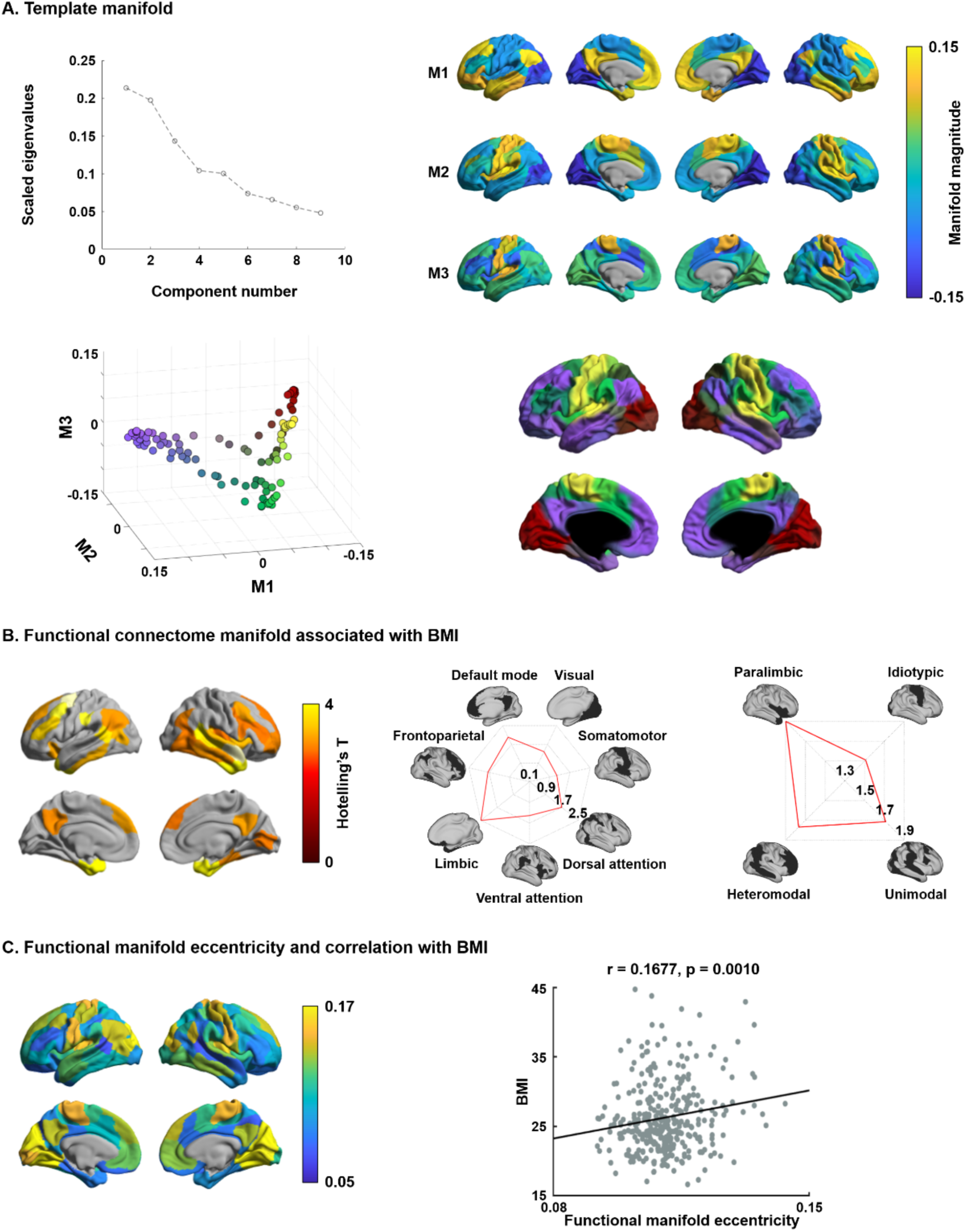
Results based on the Schaefer 100 atlas. For details, see *Fig. 1 and 2*.

**Fig. S6.**
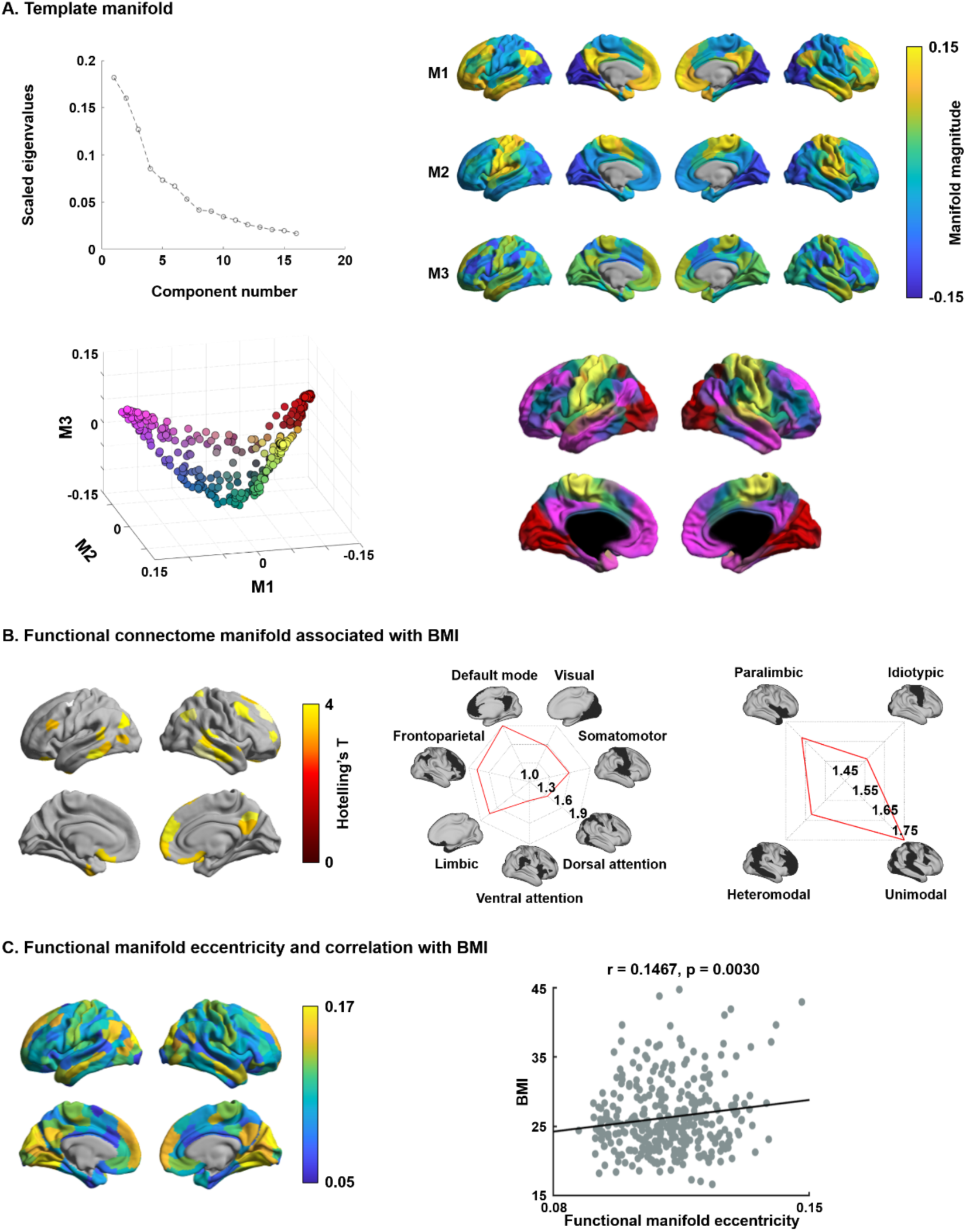
Results based on the Schaefer 300 atlas. For details, see *Fig. 1 and 2*.

**Fig. S7.**
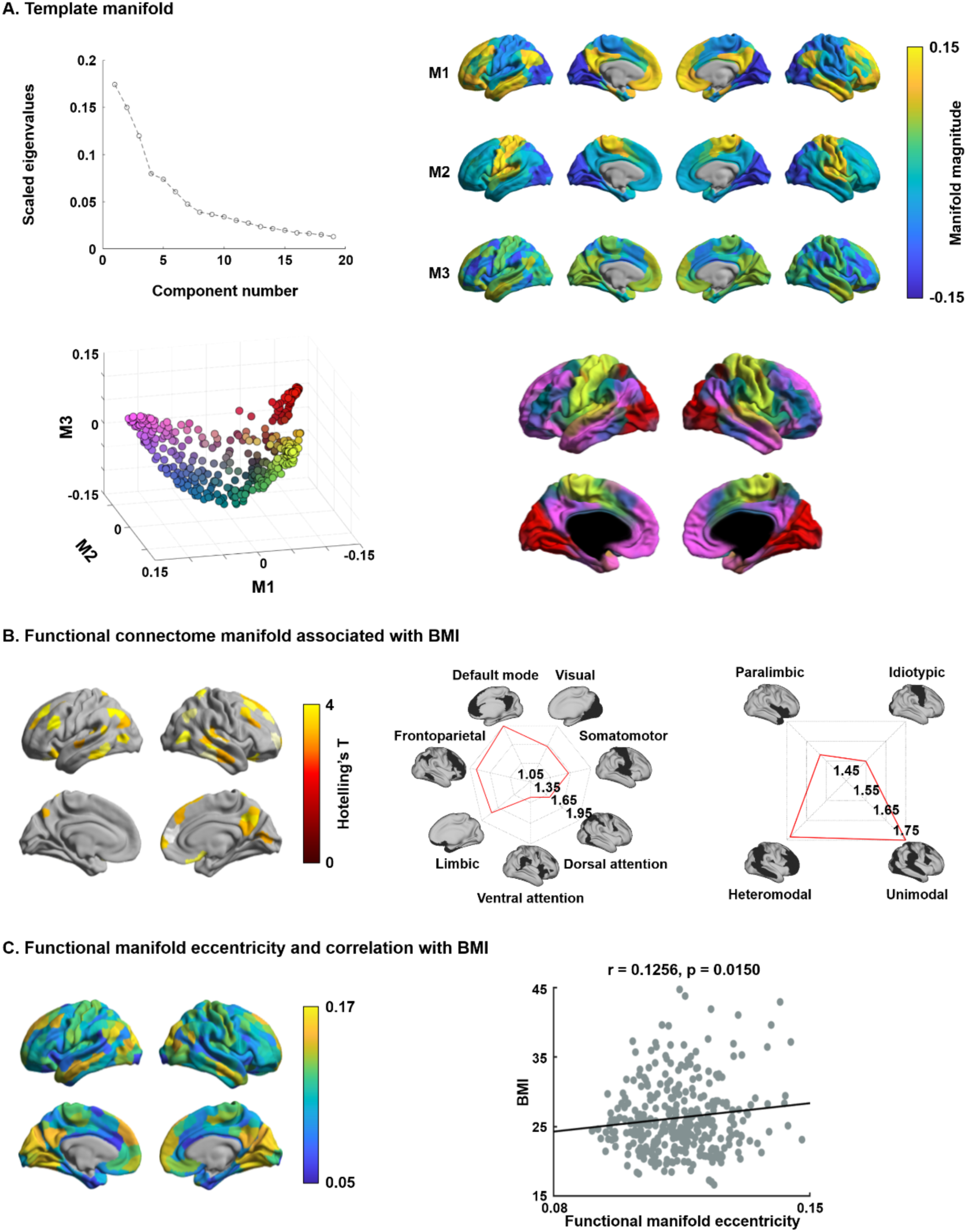
Results based on the Schaefer 400 atlas. For details, see *Fig. 1 and 2*.

**Fig. S8.**
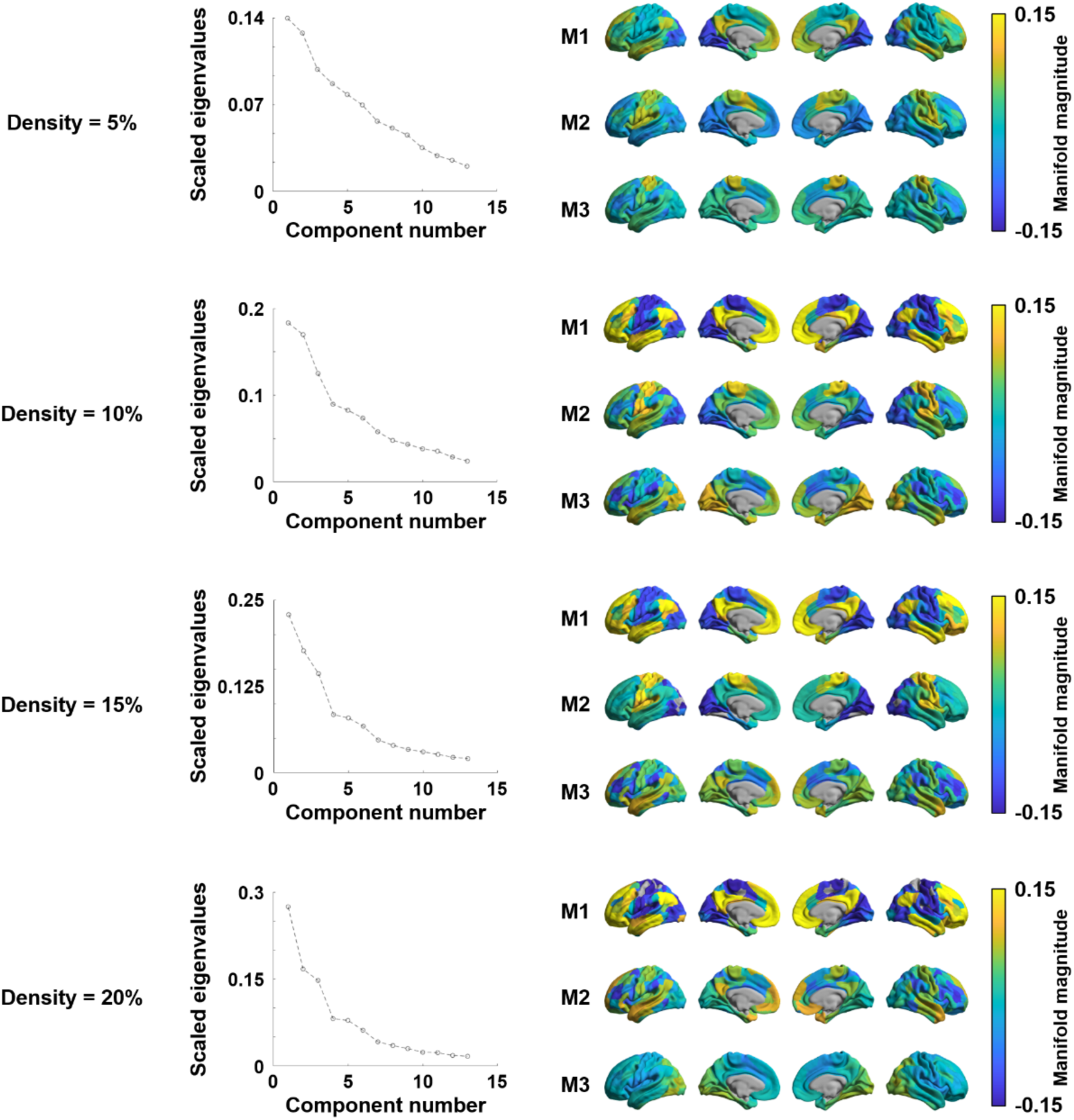
Functional manifolds with different connectome densities. Connectome density from 5% to 20% with 5% interval was applied.

**Fig. S9.**
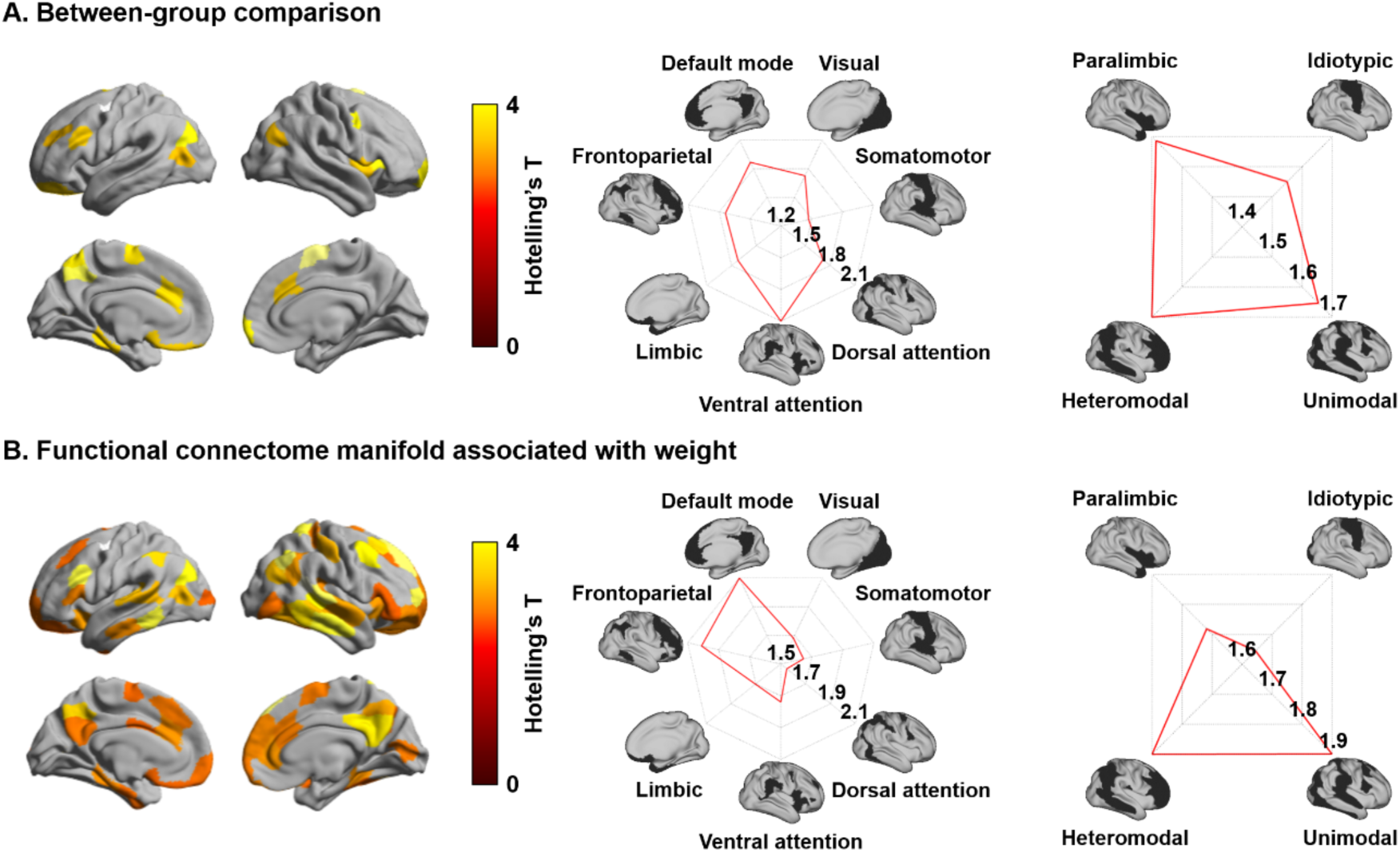
Results based on different approaches. **(A)** The t-statistics of the identified regions that showed significant between-group differences in functional connectome manifolds between individuals with healthy (18.5 ≤ BMI < 25) and non-healthy weight (BMI ≥ 25). **(B)** Those for the multivariate association analyses between functional connectome manifolds and weight.

**Fig. S10.**
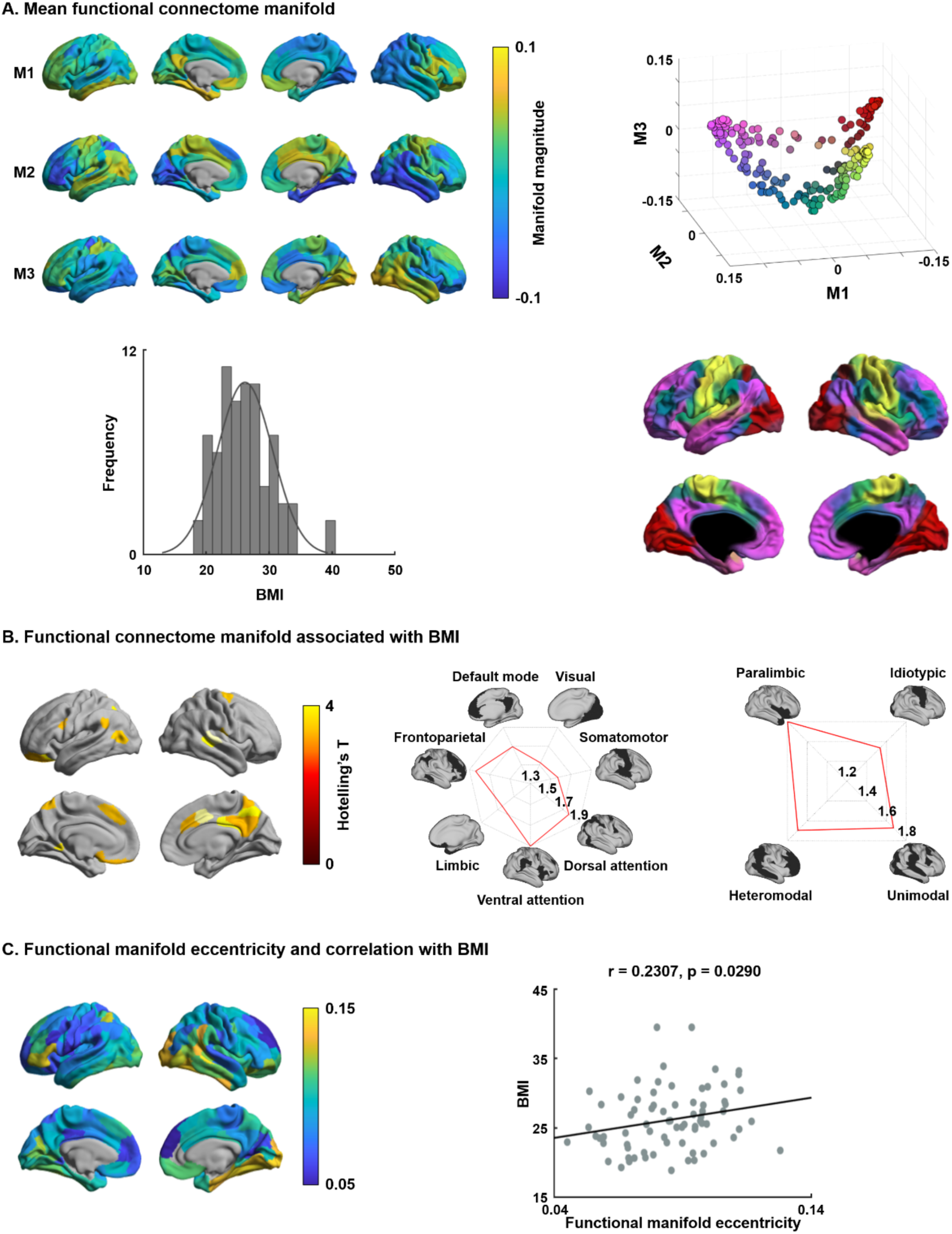
The results derived using HCP validation data. For details, see *Fig. 1 and 2*.

**Fig. S11.**
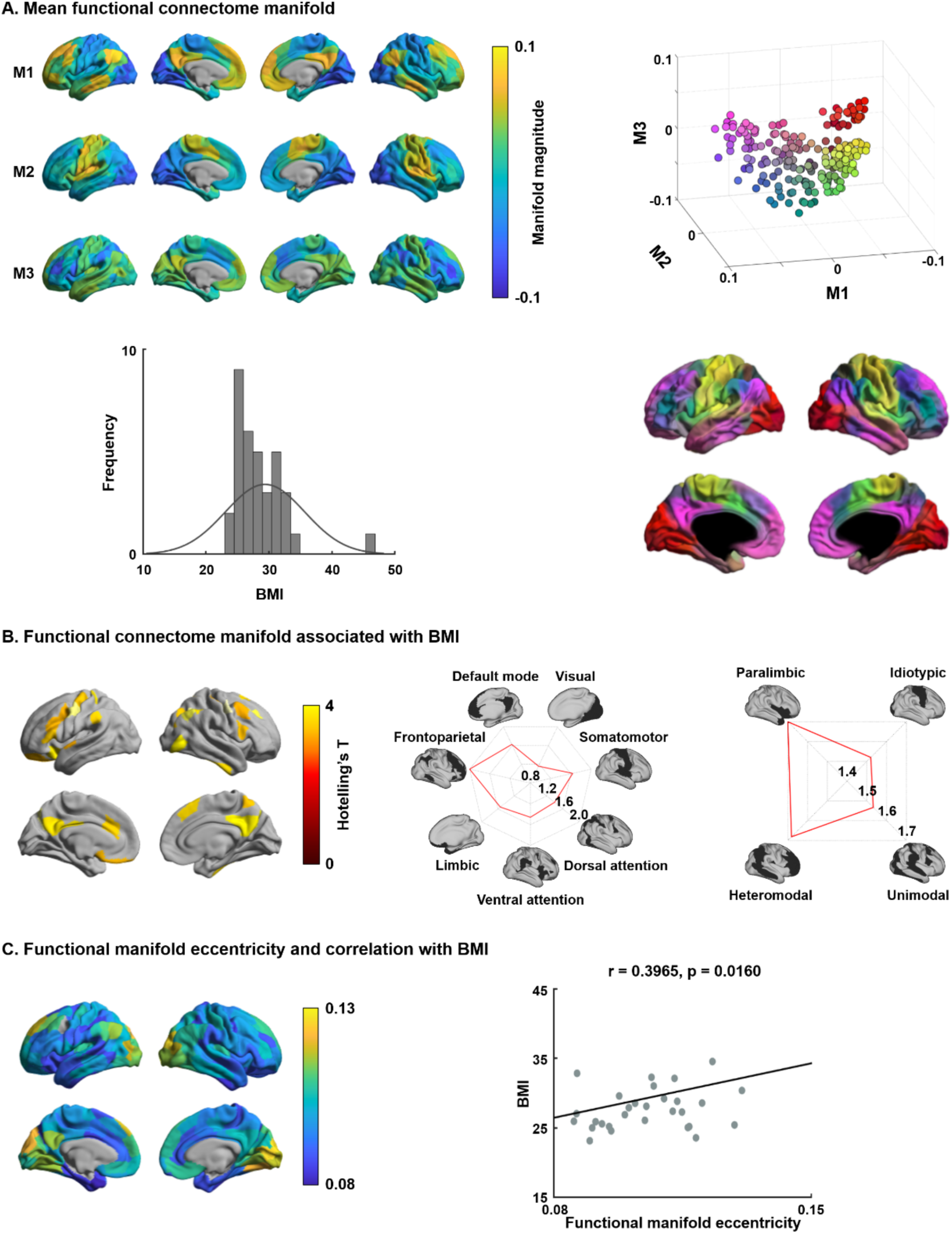
The results derived using an independent data from SVH. For details, see *Fig. 1 and 2*.

**Data S1 | Significant gene lists correlated with functional connectome manifolds associated with BMI.** Gene symbol with name and t-statistic as well as FDR corrected p-value are reported in the Supplementary Data file (Supplementary_Data1.xlsx).

